# Probiotic lactic acid bacteria induce regulatory functions in γδ T cells

**DOI:** 10.1101/2025.11.21.689796

**Authors:** Alato Okuno, Suguru Saito

## Abstract

While the modulatory effects of probiotic LAB on conventional T cells are well documented, its effects on unconventional T cells, including gamma-delta (γδ) T cells, have not yet been fully characterized. In this study, we show that probiotic LAB has the capacity to generate regulatory γδ T cells. Probiotic LAB stimulation significantly increased interleukin (IL)-10, but not IFN-γ or IL-17A, in γδ T cells compared to stimulation with methicillin-resistant Staphylococcus aureus (MRSA). This regulatory function subsequently attenuated IL-8 production in intestinal epithelial cells. Thus, probiotic LAB can induce regulatory functions in γδ T cells, which may contribute to the maintenance of intestinal health.

---

It is well documented that probiotic LAB can modify the function of immune cells [1]. This immunomodulatory effect is frequently described in myeloid lineage cells, such as dendritic cells (DCs) and macrophages, both of which are antigen-presenting cells (APCs) [2, 3]. The major mechanism underlying the functional modification induced by probiotic LAB is based on enhanced cellular activation through pattern recognition receptors (PRRs), such as Toll-like receptors (TLRs), C-type lectin receptors (CLRs) and NOD-like receptors (NLRs), similar to the recognition of pathogenic bacteria [4, 5]. This stimulatory signal ultimately regulates gene expression, resulting in enhanced cellular activities such as cytokine production, migration, antigen uptake and processing, antigen presentation, and metabolism in the APCs [1, 2, 3, 6, 7]. These enhanced functions of APCs ultimately can activate T cells; therefore, the immunomodulatory effects of probiotic LAB can alter both innate and adaptive immunity.

While several studies have documented the role of probiotic LAB in immune activation, including our previous works, the cell types examined remain limited. To expand our understanding of how probiotic LAB influence the entire immune system, further investigation is required across a broader range of immune cell subsets. We have focused on the potential immunomodulatory effects of probiotic LAB on innate T cells, such as γδ T cells, natural killer T (NKT) cells, and mucosal-associated invariant T (MAIT) cells [8-10]. Innate T cells are a distinct population from conventional thymus-derived TCRαβ T cells and are therefore considered to have a different developmental origin [8-10]. In fact, some innate T cells can self-proliferate and acquire functional polarization within tissues and organs [8, 11]. Additionally, these cells share characteristics with innate immune cells, including the expression of various PRRs, which are minimally expressed on conventional T cells [8-10, 12]. Thus, we hypothesized that innate T cells may be capable of responding to probiotic LAB and acquiring functional modification.

Here, we show that probiotic LAB have the capacity to induce regulatory function in γδ T cells. To investigate whether probiotic LAB modify γδ T cell function, we treated human peripheral blood mononuclear cells (PBMCs) with heat-killed (HK) LAB or pathogenic bacteria, methicillin-resistant *Staphylococcus aureus* (MRSA). We used *Lactococcus lactis* subsp. *cremoris* C60 and *Lactococcus lactis* subsp. *lactis* L8 as representative strains that have already been characterized for their probiotic functions [1-3, 6, 7, 13]. PBMCs (1.0×10^7^/mL, obtained from ZenBio, Durham, NC, USA) were incubated in RPMI complete medium (supplemented with 10% Fetal Bovine Serum (FBS), 100 U/mL penicillin, and 100 μg/mL streptomycin) in the presence of control (PBS), HK-C60 (5.0×10^8^ CFU/mL), HK-L8 (5.0×10^8^ CFU/mL), or MRSA (3.0×10^8^ CFU/mL) at 37°C for 24 h. The samples were then subjected to flow cytometry analysis. All fluorochrome-conjugated monoclonal antibodies (mAbs) used in this study were purchased from BioLegend (San Diego, CA, USA).

The frequency of γδ T cells within the CD45^+^ population did not change across the treatment conditions (Figure 1A). However, cytokine production showed dramatic differences between probiotic LAB- and MRSA-treated cells. The frequencies of interferon-gamma (IFN-γ) or interleukin (IL)-17A producing γδ T cells were significantly increased in HK-C60- or HK-L8-treated groups compared to control, and were further elevated in MRSA-treated cells (Figure 1C–E). Interestingly, probiotic LAB markedly increased IL-10 production in γδ T cells, with a slightly stronger effect in HK-C60-treated cells than in HK-L8-treated cells. In contrast, MRSA did not induce IL-10 production, and its levels were comparable to control (Figure 1C, F). These cytokine secretion profiles were preserved in isolated γδ T cells stimulated with HK-LAB or HK-MRSA (Supplemental Figure 1). This indicates that γδ T cells can directly recognize bacterial stimuli and become activated without assistance from other immune cells, including antigen-presenting cells (APCs).

**Figure 1.**
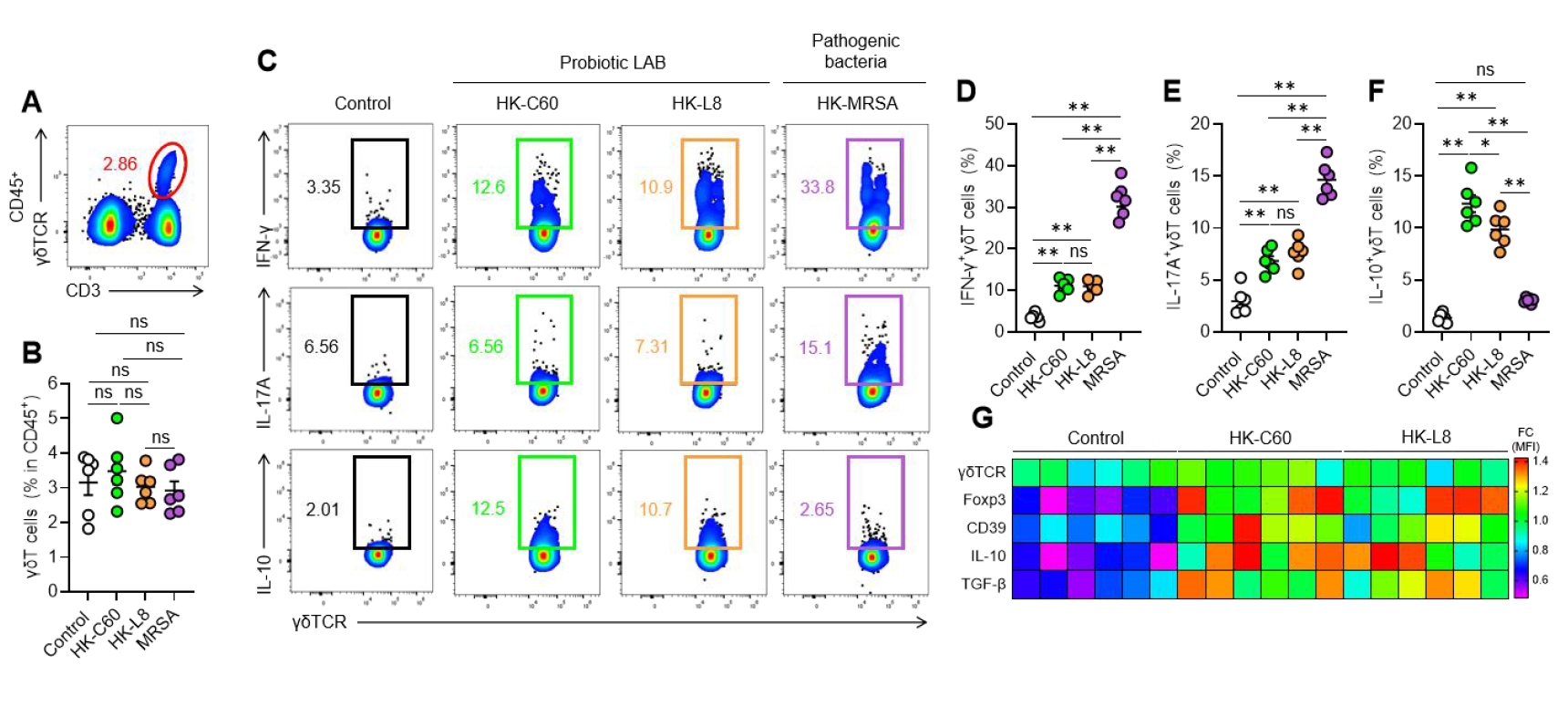
Probiotic LAB induces polarization of γδT cells into regulatory function. Human PBMCs were cultured with control, HK-C60, HK-L8 or HK-MRSA at 37C for 24 h. The sample was analyzed by flow cytometry. A-B) The frequency of γδT cells in CD45+ total leukocytes. Representative plot (A) and cumulative percentages (B) of γδT cells are shown. C) Representative plot of Cytokine producing population in γδT cells. Cumulative percentage of IFN-γ^+^(D), IL-17A^+^(E) or IL-10^+^(F) γδT cells are shown. G) The expression of regulatory γδT markers. The data are shown as mean ± SEM of six samples from two independent experiments. One-way ANOVA was used to analyze data for significant differences. Values of \**p* < 0.01 and \*\**p* < 0.001 were regarded as significant. ns: not significant.

Since increased IL-10 production is a key feature of regulatory (anti-inflammatory) γδ T cells, we analyzed markers associated with regulatory γδ T cell identity [14, 15, 16]. All tested molecules, including forkhead box P3 (Foxp3), CD39, IL-10, and transforming growth factor-β (TGF-β), were upregulated in probiotic LAB-treated γδ T cells compared to control (Figure 1G). Thus, probiotic LAB are capable of inducing regulatory function in γδ T cells.

Next, we proceeded to investigate whether probiotic LAB-stimulated γδ T cells have a suppressive effect. To do so, we employed a co-culture system using PBMC-isolated γδ T cells and Caco-2 cells, a human intestinal epithelial cell line (colorectal adenocarcinoma) [17]. Caco-2 cells were seeded in plates and used for experiments after reaching more than 80% confluency. PBMCs (1.0×10/mL) were treated with control (PBS), HK-C60 (5.0×10^8^CFU/mL), or HK-L8 (5.0×10^8^ CFU/mL) in RPMI complete medium at 37°C for 24 h, and γδ T cells were then isolated from the cultures using the EasySep™ Human Gamma/Delta T Cell Isolation Kit (STEMCELL Technologies, Vancouver, BC, Canada). Caco-2 cells were primed with lipopolysaccharide (LPS; 100 ng/mL, E. coli O55:B5, Thermo Fisher Scientific, Waltham, MA, USA) at 37°C for 6 h and then washed with PBS. The LPS-primed Caco-2 cells and pre-treated γδ T cells were co-cultured in 24-well plates without direct contact using transwell inserts. The ratios between Caco-2 cells and γδ T cells were tested at 1:0.1, 1:0.25, and 1:0.5. After incubation at 37°C for 16 h, IL-8 concentration in the conditioned media was measured by ELISA under each culture condition (Figure 2A).

**Figure 2.**
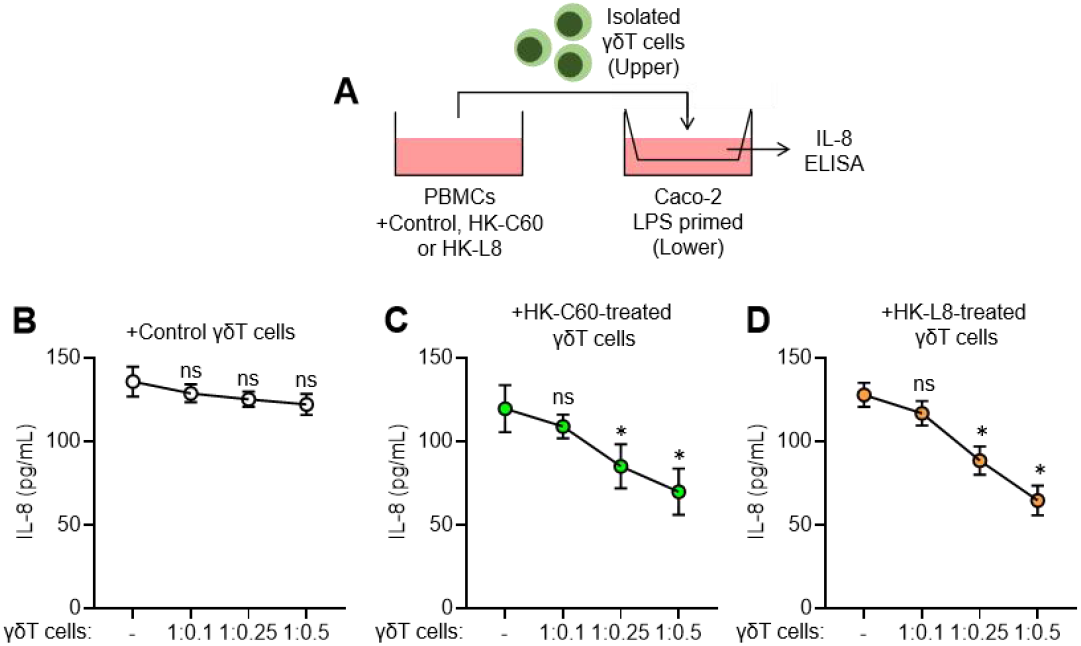
Suppressive effect of probiotic LAB-stimulated γδT cells. A) Experimental design of *in vitro* suppressive assay. Caco-2 cells were primed with LPS at 37C for 6 h. PBMCs were pre-treated with control (PBS), HK-C60 or HK-L8 at 37C for 24 h. The γδT cells were isolated by magnetic sorting, then transferred to upper chamber of trans well placed on Caco-2 culture. After co-incubation at 37C for 16 h, the conditioned media was harvested and subjected to measuring IL-8 concentration by ELISA. IL-8 concentration in the Csco-2 culture with control-treated γδT cells (B), HK-C60-treated γδT cells (C), HK-L8-treated γδT cells (D). The data are shown as mean ± SD of six samples from two independent experiments. Student t-test was used to analyze data for significant differences. Value of \**p* < 0.001 was regarded as significant. ns: not significant.

As expected, LPS stimulation induced IL-8 production in Caco-2 cells. The IL-8 level was not suppressed by control γδ T cells pre-treated with PBS (Figure 2B). In contrast, γδ T cells pre-treated with HK-C60 or HK-L8, which were expected to have regulatory function, showed a suppressive effect against IL-8 production in LPS-primed Caco-2 cells. This suppressive effect became significant at ratios equal to or greater than 1:0.25 (Figure 2C, D). Taken together, the regulatory function of probiotic LAB-stimulated γδ T cells enables attenuation of inflammatory responses in intestinal epithelial cells.

Innate T cells, including γδ T cells, are relatively enriched in the intestinal mucosa, suggesting that orally administered probiotic LAB can interact with these cells in the gut environment [8, 14, 15]. Functional polarization toward a regulatory phenotype upon probiotic LAB stimulation may increase the contribution of γδ T cells to anti-inflammatory regulation in the intestine. Indeed, regulatory γδ T cells have been proposed to play an important anti-inflammatory role in the gut [14-16]. Other anti-inflammatory T cell subsets, such as regulatory T cells (Treg) and type-I regulatory T cells (Tr1), are also known to be important in maintaining intestinal immune tolerance [18, 19]. However, regulatory γδ T cells may have a unique advantage in intestinal immune control. Unlike Treg and Tr1 cells, whose induction requires antigen presentation and cytokine-driven signaling, γδ T cells may respond directly to probiotic LAB through PRRs, similar to myeloid cells [8, 12]. This “self-reprogramming” ability may allow γδ T cells to adopt regulatory function more rapidly than conventional T cells during intestinal inflammation.

A previous study also reported that probiotic LAB can increase regulatory γδ T cell populations [20], although the direct suppressive function was not clearly demonstrated. In contrast, our study shows direct suppression of inflammatory cytokine production in intestinal epithelial cells. Additionally, we characterized both cytokine secretion and regulatory surface markers in γδ T cells following probiotic LAB exposure, distinguishing our findings from earlier reports. However, the key molecular factors responsible for this anti-inflammatory activity remain unclear. We hypothesize that the significantly increased IL-10 may play a central role in the regulatory function of probiotic LAB-stimulated γδ T cells. The involvement of cell-to-cell contact mechanisms, including immune checkpoint molecules (ICMs), also warrants investigation [21]. Moreover, the suppressive function of probiotic LAB-stimulated γδ T cells may extend to other inflammatory immune cells, such as Th1 and Th17 cells, further contributing to intestinal immune homeostasis.

This study also raises important questions about the receptor(s) responsible for triggering regulatory polarization in γδ T cells. Because γδ T cells express multiple PRRs, these receptors represent potential candidates. Downstream transcription factors, including Interferon Regulatory Factors (IRFs), Nuclear factor-kappa B (NF-κB), and Activator Protein-1 (AP-1), should also be examined for activation upon probiotic LAB stimulation [8, 22, 23]. Based on our current findings, we are also interested in whether probiotic LAB influence other innate T cells, such as NKT cells and MAIT cells, as well as innate lymphoid cells (ILCs), all of which may contribute to anti-inflammatory regulation in the intestine [24, 25]. Although these cell types differ in phenotype from γδ T cells, we hypothesize that these non-myeloid innate immune cells may also respond to probiotic LAB and gain functional modulation.

Overall, our study provides key insights into the role of probiotic LAB in generating regulatory γδ T cells and contributes to a broader understanding of how probiotic LAB shape immune homeostasis.

## Supporting information

Supplemental Meterials

## Author Contributions

Conceptualization, S.S.; Methodology, S.S.; Experiments, A.O. and S.S.; Data Analysis, A.O. and S.S.; Resources, A.O. and S.S.; Discussion, A.O. and S.S; Writing Manuscript, A.O. and S.S.; supervision, S.S.; Project Administration, A.O. and S.S.; Funding Acquisition, A.O. and S.S. All authors have read and agreed to the published version of the manuscript.

## Acknowledgement

We thank the National Agriculture and Food Research Organization (NARO) for providing *Lactococcus lactis* subsp. *cremoris* C60. We also thank LAVIEPRE Co., Ltd. for providing *Latococcus lactis* subsp. *lactis* L8.

