## Supplemental Meterials for "Probiotic lactic acid bacteria induce regulatory functions in γδ T cells"


---

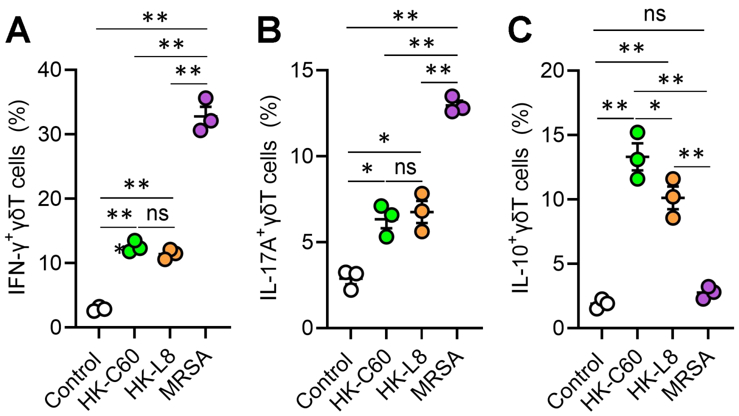


**Supplemental Figure 1. Cytokine production of stimulated γδT cells**

The γδT cells ($1.0\times10^7$/mL) were isolated from PBMCs, then incubated with control (PBS), HK-C60 ($5.0\times10^8$ CFU/mL), HK-L8 ($5.0\times10^8$ CFU/mL) or MRSA ($3.0\times10^8$ CFU/mL) at 37°C for 24 h. The cytokine productions were analyzed by flow cytometry. The frequencies of IFN-$\gamma^+$ γδT cells (A), IL-17A$^+$ γδT cells (B) and IL-10$^+$ γδT cells (C) were shown, respectively. The data are shown as mean ± SEM of three samples. One-way ANOVA was used to analyze the data for significant differences. Values of $*p < 0.01$ and $**p < 0.001$ were regarded as significant. ns: not significant.
